# Optimal distance metrics for single-cell RNA-seq populations

**DOI:** 10.1101/2023.12.26.572833

**Authors:** Yuge Ji, Tessa D. Green, Stefan Peidli, Mojtaba Bahrami, Meiqi Liu, Luke Zappia, Karin Hrovatin, Chris Sander, Fabian J. Theis

## Abstract

In single-cell data workflows and modeling, distance metrics are commonly used in loss functions, model evaluation, and subpopulation analysis. However, these metrics behave differently depending on the source of variation, conditions and subpopulations in single-cell expression profiles due to data sparsity and high dimensionality. Thus, the metrics used for downstream tasks in this domain should be carefully selected. We establish a set of benchmarks with three evaluation measures, capturing desirable facets of absolute and relative distance behavior. Based on seven datasets using perturbation as ground truth, we evaluated 16 distance metrics applied to scRNA-seq data and demonstrated their application to three use cases. We find that linear metrics such as mean squared error (MSE) performed best across our three evaluation criteria. Therefore, we recommend the use of MSE for comparing single-cell RNA-seq populations and evaluating gene expression prediction models.

## Introduction

In systems biology, profiling the transcriptomes of individual cells by single-cell RNA sequencing (scRNA-seq) has become a common tool for gaining insight into the function of complex biological systems. When analyzing or modeling scRNA-seq data, we rely on distance metrics to capture relevant differences in the data. For instance, we might use the t-statistic to identify differentially expressed genes and mean squared error in the reconstruction loss of a generative model. These distance metrics make assumptions about the data which may not always hold. If the assumptions of the metric do not match the specific data distribution, the resulting analysis or model is likely to suffer. Therefore, it is essential to pick a metric that is suitable for the data characteristics.

One application area is generative modeling of transcriptome data, where computational models are trained to reconstruct a cellular response on the transcriptome level. These models are usually evaluated based on how close their predicted gene expression profile is to an experimentally determined ground truth response. Ultimately, generative models are used to predict the changes of cells transcriptomes in response to time, a drug treatment or genetic alteration. There is currently no consensus in the literature on which distance metric to use for evaluating these models.

Many generative models^1–3^, as well as linear^4^, graph-based^5,6^, dynamic^7,8^ and optimal transport-based models^9,10^ output predicted expression data that aims to accurately reflect the state of cells after a specific intervention^11^. Different publications have used the coefficient of determination (R^2^)^1,12^, maximum mean discrepancy (MMD)^13^, Spearman correlation^13^, Wasserstein^14^ and mean absolute error (MAE)^15^ for evaluation, each with its own set of assumptions and limitations. Despite the relevance of developing generative models, for example in reducing the search space of drug or genetic screens, there is a lack of comparative assessment of these distance metrics for scRNA-seq generative model evaluation. As such, a thorough investigation of distance metrics with a focus on biological interventions is timely for improving model development.

Here, we provide a reusable framework for evaluating distance metrics for single-cell gene expression data. To mimic how distance metrics would be used in model evaluation or dataset analysis, we quantify their sensitivity and robustness when identifying differences between populations (Figure 1a,b). We also quantify each metric’s ability to capture variation that is expected based on prior knowledge (Figure 1c). This work uses ground truths such as developmental time and small molecule perturbation, contrasting with previous work using cell type labels^49^. To ensure that our results are broadly applicable, we devise three evaluation metrics—control rank percentile (CRP), robustness, and biological reproducibility (BioRep)—and apply our analyses to 16 distances across seven scRNA-seq datasets (Figure 1d). To assist users in deciding on a distance metric, we identify common trends in distance metric behavior, segregating the metrics into three categories based on their performance under various technical effects on scRNA-seq data. We show use cases in model training, model evaluation, and cell type analysis to demonstrate that the choice of distance metric matters for downstream tasks.

**Figure 1:**
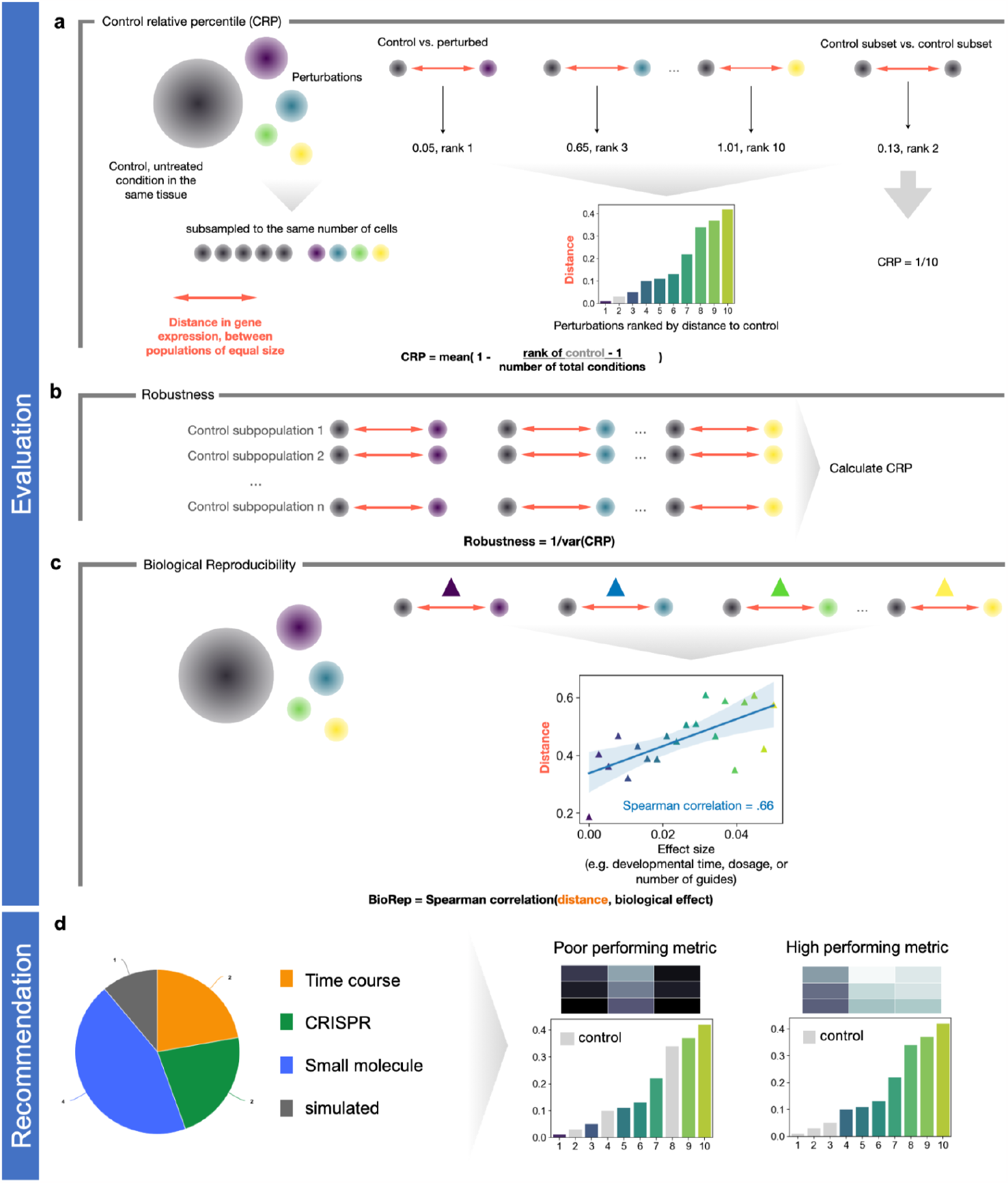
Evaluation measures used for cell population distance metrics evaluation. (a) *Control rank percentile (CRP)* measures how close random samples of the control cells are to each other relative to how close they are to perturbed populations. Distances were measured between control pairs and control-perturbation pairs, ordered by magnitude, and then the relative rank of the control pairs was used to assess if the metric can distinguish between perturbed and control populations. (b) *Robustness* measures how similar control–perturbation distances are when calculated for different random samples of the control cells. (c) *Biological reproducibility (BioRep)* measures how well the measured distance metrics correspond to biological ground truth based on a biological covariate. (d) Recommendations were formulated from multiple types of experimental and simulated datasets. High-performing metrics have control populations that were close to each other and had a high correlation with known covariates.

## Methods

### Distances

We selected distance metrics based on common usage in genomics analysis and machine learning, including those used previously for model evaluation, as loss functions in model training, or for data analysis. We tested classically used distances (Euclidean, cosine distance, mean absolute error (MAE), mean squared error (MSE), linear maximum mean discrepancy (linear MMD)^16^, Wasserstein^17,18^), in addition to some statistics commonly used in biological analysis (t-statistic, Pearson correlation, Spearman correlation, Kendall tau distance, coefficient of determination (R^2^), energy distance (E-distance)^19^), and previously unexplored ones (Kolmogorov-Smirnov test, symmetric Kullback-Leibler divergence, linear classification probability). After scaling of the metric, linear MMD, MSE, and Euclidean are mathematically identical, but we included all three implementations for completeness. Note that E-distance was computed directly on genes, which is different from prior use where it was computed on PCA embeddings.

Metrics that are not formally distances, such as Pearson correlation, were converted to return positive values (Table 1). Non-distributional distances were computed between the centroids of the two conditions, in accordance with conventional pipelines for single-cell omic data^20,21^. Defining equations can be found in the Supplementary Methods.

**Table 1:** Summary of evaluated metrics. The four distance axioms referred to are I. Definiteness, II. Symmetry, III. Triangle inequality, and IV. Positivity. Representative distances for type I and type II are bolded. Conversion to distance ‘-’ when no conversion needed.

| Metric | Abbrev. | Conversion to distance | Assumptions | Differentiable | Axioms satisfied | Type |
| --- | --- | --- | --- | --- | --- | --- |
| **Mean squared error** | MSE | - | Gaussian | yes | yes | Ia |
| Maximum mean discrepancy | MMD | - | Depends on kernel | yes | yes | Ia |
| Euclidean distance | Euclidean | - | None | yes | yes | Ia |
| Energy distance | E-distance | - | Gaussian | yes | yes | Ia |
| Kolmogorov–Smirnov test | KS test | - | None | no | yes | Ia |
| Mean absolute error | MAE | - | Laplacian | yes* | yes | Ib |
| Two-sided T-test statistic | T-test | - | Gaussians with equal variance | yes | violates III | Ib |
| **Cosine distance** | cosine | - | None | yes | yes | II |
| Pearson correlation coefficient | Pearson | 1 - r | Linearity, Gaussian | yes | violates III | II |
| Coefficient of determination | R<sup>2</sup> | 1 - R<sup>2</sup> | None | yes | violates II, III | II |
| Classifier control probability | classifier proba | 1 - control probability | None | no | violates II, III | II |
| Kendall tau distance | Kendall tau | - | Ordinal variables | no | yes | III |
| Spearman correlation coefficient | Spearman | 1 - rho | Ordinal variables | no | violates III | III |
| Wasserstein distance | Wasserstein | - | Corresponding populations | yes | violates I | III |
| Symmetric Kullback-Leibler divergence | symmetric KL | Gaussian for non-discrete formulation | X, Y occupy same distribution space, Gaussian | yes | violates III | III |
| Classifier class projection | classifier_cp | 1 - control probability | None | no | violates II, III | - |

### Datasets

We used one simulated and six public scRNA-seq datasets for evaluation (Supplementary Table 1).

Simulated data was generated using Splatter^22^ (Supplementary Methods). The control condition was modeled as a heterogeneous population. 32 perturbed conditions were generated using all pairwise combinations of eight values representing different probabilities that a gene is differentially expressed (‘DEProb’) and four values representing varying magnitudes of corresponding differential expression (‘FacLoc’).

Six public datasets were selected representing common multi-condition experimental setups (Figure 2a), such as time-course^9,23^, CRISPR perturbation^24,25^, or small molecule treatment^26,27^. Five out of the six contain covariates were expected to correlate with effect strength per condition, such as increasing time, multiple dosages, or more than 1 guide RNA (gRNA). To ensure perturbation effect was not confounded with other biological factors such as cell type, each dataset was filtered to a single cell-type with the most cells across conditions. The exceptions were Sci-plex3, in which all three contained cell types were sufficiently abundant and therefore used separately, and Santinha et al. where all cell types were included. Datasets varied greatly in how strongly the labeled control condition and intervention transcriptomes differed (Figure 2a). For instance, datasets from large-scale screens based on CRISPR or small molecules contained many conditions which likely did not cause a significant change from control.

**Figure 2:**
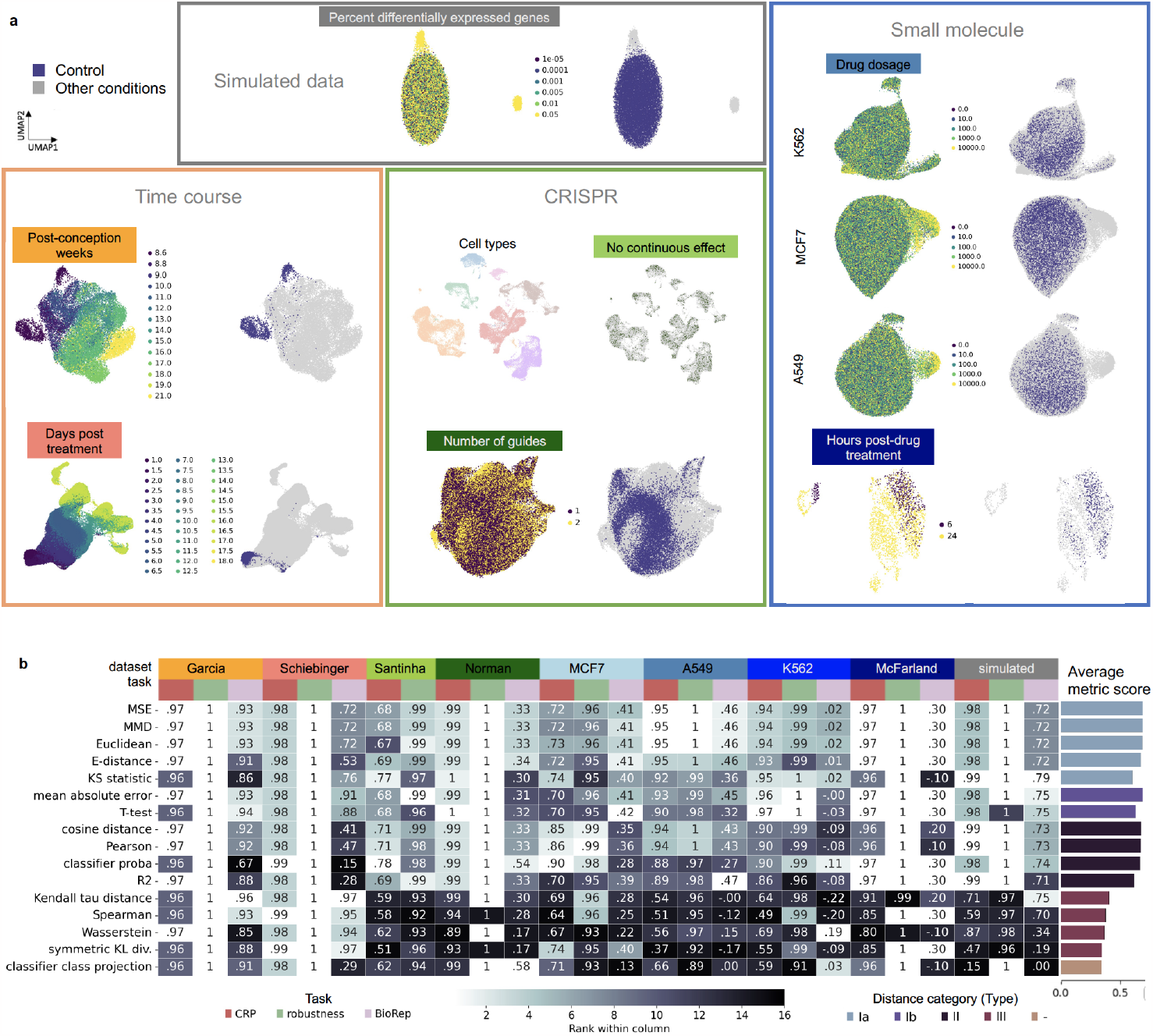
Description of intervention datasets and distance metric performance results. (a) Categories used for evaluation. For each dataset the UMAP on the left is colored by the biological covariate used when calculating BioRep metric; the UMAP on the right highlights in blue the control population used when calculating CRP. The three labeled K562, MCF7, and A549 are different cell lines from a single experiment (Sci-plex3). (b) Overview of distance metric performance for each dataset and metric evaluation method, colored by rank across the evaluated metrics. The mean score for each metric is shown on the right, colored by the category each distance was assigned to (referred to as “type” below). Distances of the same type have similar absolute values. Type I distances performed better overall across tasks and datasets.

All benchmarks were computed using unscaled, log-normalized counts, with conditions subsampled to the same number of cells and 2000 highly variable genes calculated jointly across all perturbation conditions, including control, using scanpy^28^ with default parameters (Supplementary Methods). We also evaluated distances on raw count data using the same filtering and highly variable genes, but without log-normalization. Performance is reported only for log-normalized data, as performance on raw count data was comparable (Supplementary Figure 2d).

### Distance evaluation

We defined desirable properties of a good distance metric as follows. First, given prior knowledge, it should assign high distance values between cell populations with expected biological differences. Second, it should do so robustly, i.e. with minimal sensitivity to noise inherent to scRNA-seq. Thus, we reasoned that distances should perform well in differentiating untreated or baseline conditions (“control”) from conditions where an intervention has been applied (“perturbed”).

To this end, we defined an evaluation measure **“control relative percentile” (CRP)** (Figure 1a). For each dataset, we randomly sampled five disjoint sets of cells of equal size (n=300) from the control population. We selected five sets to compute the variance for each distance while still retaining enough cells per sample. For each set of control cells, distances were computed to all other cell populations, including the perturbed populations and the other four control sets. **The CRP is defined as the percentage of perturbed conditions with a larger distance to the reference control set than the control sets to each other, averaged across five control sets**. For an ideal metric, this would equal one. We provide pseudocode for the CRP measure in the Supplementary Methods.

The primary assumption was that control cells should always be closer to each other than to perturbed cells unless the distance metric cannot distinguish between control and perturbed. It could also be true that the drug or CRISPR intervention had no effect. However, this would not lead to biases when comparing different metrics, as all distance calculations would suffer equally. A *better* distance metric would still have a smaller value, as fewer conditions would be close to control.

We defined **robustness** as the inverse of the variance of CRP, such that metrics with higher scores were more robust (Figure 1b). We reasoned that it is desirable for a metric to be as insensitive as possible to variation within a population and return similar relative distances when comparing different views of identical populations. Thus, a better distance metric should have minimal differences in CRP across the five control conditions and a high robustness score.

We define **biological reproducibility (BioRep)** to examine whether metrics correlated with biologically meaningful continuous effect such as drug dosage, developmental time, or the number of gRNAs targeting different genes. This contrasts with CRP and robustness, which only measure the binary question of whether a distance finds that perturbations had an effect. For each dataset, we calculated the Spearman correlation between each metric’s value for each perturbed condition and the corresponding biological covariate (Figure 1c). This test approximates how much a distance aligns with our expectation of effect sizes based on prior knowledge.

When comparing between metrics, values were rounded to three decimal places so that extremely similar performance would result in a tie. The final order of which metric performed best was decided by how many times a metric performed at or above average (Figure 1d).

## Results

### Our framework successfully characterizes similarities and differences between distance metrics

Most metrics performed reasonably well across all datasets. The average CRP, robustness, and BioRep across all metrics and all datasets was 0.62, 0.77, and 0.66 respectively. This indicates that on average metrics could robustly distinguish 62% of interventions from control, a reasonable number given that many interventions are not expected to have an effect in large screens. We can also see that metrics were positively correlated with effect sizes, indicating that they indeed captured meaningful biological variation. This supports that CRP, robustness, and BioRep are evaluation measures which are neither too simplifying nor too difficult to compute and comprehend, making them reasonable measures of distance metric performance.

In addition, all three evaluation measures showed variation across metrics, indicating that they are able to distinguish between the performances of individual metrics. Distance metrics generally performed consistently across datasets, supporting the robustness of this measure across different biological systems, batch effects, and control population definitions. The consistent behavior across datasets adds confidence that our recommendations will apply to other scRNA-seq datasets.

To summarize and simplify the comparison of distances, we grouped them by performance and behavior (Table 1, last column). We excluded classification class projection due to extremely subpar performance (Figure 2b) and clustered the remaining 15 distances into three different categories—type I, II, and III—based on projection of performance into a PCA space (Supplementary Figure 2a). Type I distances were separated into type Ia and Ib due to slightly different behavior between the two groups; type Ib performed better on BioRep (Supplementary Figure 1c). All distances in type I behaved similarly with respect to observed metric biases (Supplementary Figure 2c).

### MSE and similar distance metrics best capture interventional effects in scRNA-seq data

One desirable metric characteristic is consistently strong performance across a variety of datasets. To make a general recommendation, we aggregated results across all three evaluation measures and datasets by averaging scaled evaluation scores (Figure 2b). MSE and the mathematically identical linear MMD performed above the median of all metrics in all datasets. Euclidean distance and MAE performed comparably, which is expected given their mathematical similarity to MSE and linear MMD. E-distance significantly underperformed in BioRep in Schiebinger et al. but somewhat outperformed in four other datasets. Altogether, these distances performed extremely similarly, justifying their grouping in type I (Table 1). We emphasize that type I metrics are closely related, and we would consider their performance to be equivalent. The marginal differences seen may become important when comparing models (Figure 3c) or cell populations with relatively small differences such as CRISPR perturbations.

**Figure 3.**
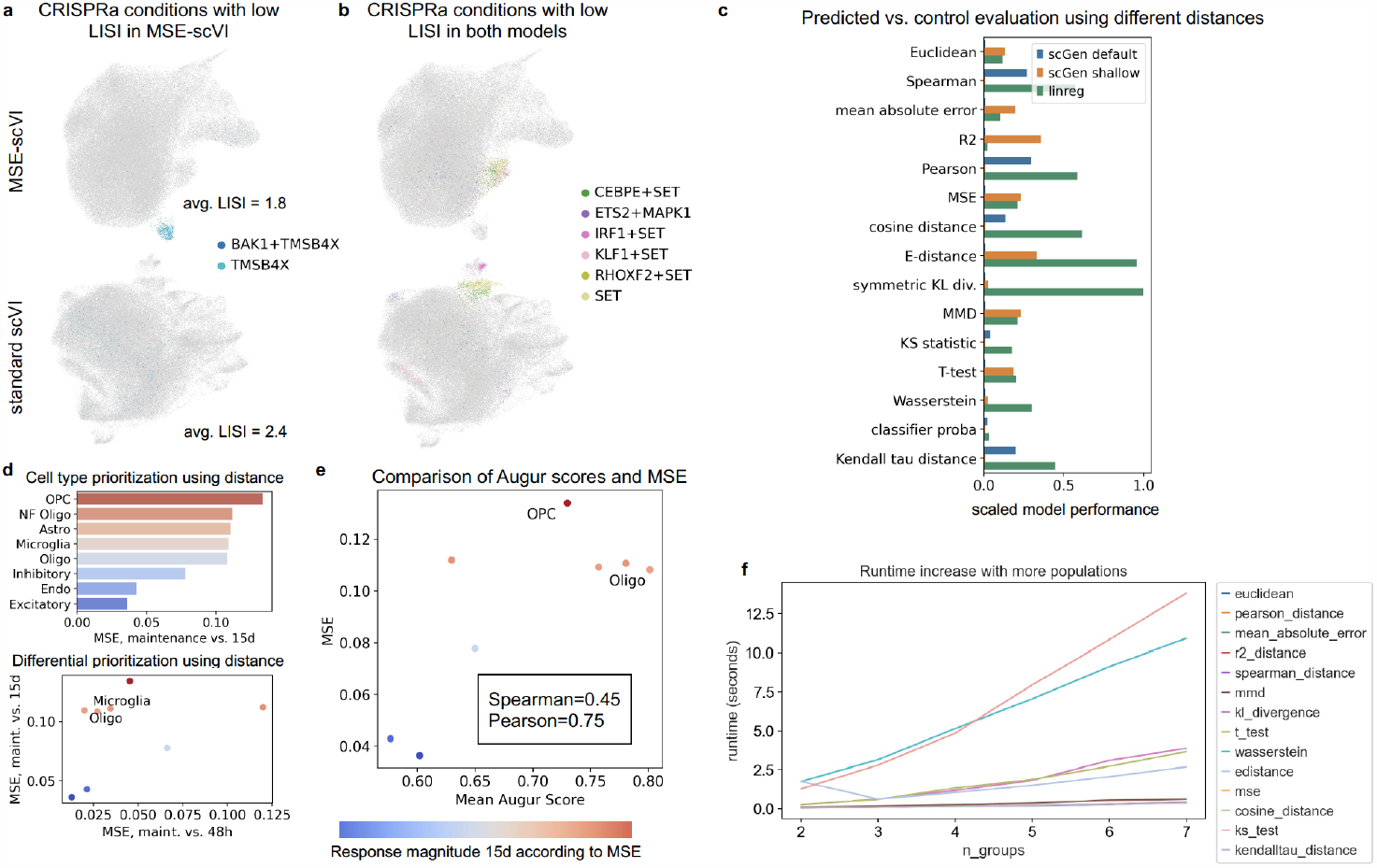
Use cases of different distance metrics and computational runtime. (a),(b) Latent representations of Norman et al. generated by two scVI-based models, colored by (a) two CRISPRa interventions which co-cluster separately from other cells in MSE-scVI but not in standard scVI and (b) six CRISPRa interventions which formed a separate cluster in the latent representations of both models. In (a), LISI score was averaged between the two interventions. In (b), the KLF1+SET CRISPRa intervention appears in a different location relative to other interventions depending on the model used. (c) Gene expression prediction performance of three different models resulted in a different winning model depending on the metric used. (d),(e) Distances quantified response relative response to perturbation across cell types. Prioritization via Augur and MSE are comparable despite dramatic differences in methodology and reduced computation time for MSE. (f) The computation time of two distances (Wasserstein and KS statistic) scaled poorly with an increasing number of conditions. Our recommended distance, MSE, increased in computational time with the number of conditions but remained fast overall.

The Type III distances (Wasserstein, symmetric KL divergence, Kendall tau distance, and Spearman correlation) performed worse than other metrics, particularly linear ones (Figure 2b, Supplementary Figures 1f). They scored an average of 0.31 and 0.51 for CRP and BioRep, respectively (Figure 2b, Supplementary Figure 1a,c) and performed worse than other distances in all datasets except Schiebinger et al. and one cell line in Sci-plex3. We believe that this instance of good performance, specifically on BioRep, is due to the Schiebinger dataset being particularly well-suited for these metrics. We hypothesize that the specific time course in Schiebinger et al. produces a smooth shift in the parameters of the underlying distribution, but not in means (Supplementary Figure 1e).

The primary distinguishing feature of type III metrics is that they capture aspects of the distribution beyond the mean and standard deviation. We hypothesize that accounting for additional moments of the distributions is detrimental to picking up intervention-related biological signal. This can be observed in low robustness scores. For instance, by using ranks, noise-induced changes in lowly expressed genes have a disproportionate effect on Spearman correlation (Supplementary Figure 2b). Noise in many lowly expressed genes may mask the true biological effects in a smaller number of significant genes in type III distances. As such, we discourage the use of type III distance metrics for evaluating the closeness of predicted cell populations to real cell populations in most standard cases.

We also note that there are differences in the rankings when examining CRP and BioRep performance separately. Among all the distances we evaluated, linear classification probability achieved the highest CRP, as expected for a classifier specifically optimizing for this task (Supplementary Methods). Among the distances without training, Pearson correlation and cosine distance performed best for this measure, scoring an average of 0.81 and 0.80 for CRP, respectively (Supplementary Figure 1a). We can expect these to be similar as Pearson correlation and cosine distance are the same for zero-centered vectors^29^. With the exception of R^2^, type II distances outperformed type III distances in CRP, and vice versa for BioRep. We hypothesize that this is because vector distances better capture absolute change, but the effects in BioRep are more a function of changes in magnitude not fully captured by vector distances.

No single metric was the top performer in more than two datasets in the BioRep evaluation. BioRep scores sometimes also differed dramatically between distances of the same type for the same dataset (Supplementary Figure 1c). This finding suggests that the distribution of gene expression changes over the course of interventions can happen in multiple ways, and may be dependent on the covariate being examined (e.g. increasing dosage or increasing time). It may also depend on the biological setting: distances from three different types were the top scorers for BioRep in each of the Sci-plex3 cell lines (Figure 2b, Supplementary Figure 1c). T-test metric performed best for MCF7, R^2^ performed best for A549, and Wasserstein performed best for K562. Thus, variation in how distributions change after intervention is potentially not just relegated to dataset-specific effects. When using distance metrics with a continuous covariate as we do to calculate BioRep, it is worthwhile to consider using a combination of several distance metrics to capture different effects.

### Representative distance metrics for data investigation

We selected MSE as the representative distance for type I and cosine distance as the representative distance for type II. MSE and cosine distance have desirable qualities in that they are already commonly employed as a loss function in regression models, are computationally fast, and are readily available in several different packages across programming languages^30–32^. They are also simple to calculate manually and interpret. Users can use these representative distances on their own data to determine if there is a category of distances that is best suited to their biological question or dataset qualities. Users can also feel confident that the other distances in the category will behave similarly to the representative distance (Figure 2b, Supplementary Figure 1a,b,c).

We do not identify a representative distance metric for type III distances because they displayed highly inconsistent BioRep scores across datasets (Supplementary Figure 1c), suggesting that they capture different qualities about what distributional shifts occur when going from control to perturbed. In the most common data analysis scenario, where dataset characteristics are not known a priori, we recommend starting with type I metrics in perturbation modeling and analysis.

### Use case 1: what a model learns changes depending on metric used

Since the absolute scores of the above evaluation measures were fairly similar across metrics (Supplementary Figure 1a,b,c), the question arises if changing the loss function of a model from one metric to another would make a substantive difference or improvement in what is being learned. To test this, we modified the loss of a generative probabilistic model, scVI^3^, and trained it on our CRISPRa dataset^24^, where no metric performed conclusively better in BioRep. We replaced the negative binomial loss, evaluated between the posterior distribution and real cells, with non-distributional MSE loss between predicted cells and real cells (Supplementary Methods). We trained with otherwise default parameters and plotted the learned latent representation of the dataset, as in a typical workflow with scVI and other generative models^20^ (Figure 3a,b). We compared this latent representation with one from a standard scVI model, using Local Inverse Simpson’s Index (LISI)^33,34^ to identify CRISPRa treatments that clustered together (Supplementary Methods). Although we found that CRISPRa treatments overall clustered less in our MSE-scVI (Supplementary Figure 3), our model also identified a new cell cluster of cells with CRISPRa activation of TMSB4X (Figure 3a).

To see if the models learned not only new but different relationships between CRISPRa conditions, we also examined the six CRISPRa treatments with low LISI scores in both models. We found that KLF1+SET was more similar to the remaining CRISPRa treatments containing SET in MSE-scVI than in standard scVI (Figure 3b). Figure 2b of Norman et al. supports this; there, KLF1+SET is clustered with other combinations including SET. Taken together, this indicates that changing the loss function of a model changes the latent space significantly, and different loss functions may provide perspective on different perturbations.

### Use case 2: model evaluation changes depending on metric used

Now that we have demonstrated that distance metrics differ in their ability to capture important variation in scRNA-seq data, we illustrate an example in which this difference affects which perturbation effect a model captures. We trained two versions of scGen^1^ with different hyperparameters, which we call “scGen default” and “scGen shallow”, using the default or a lower number of neural network layers. We compared the scGen models against linear regression as a simple baseline (Supplementary Methods). We applied the models to PBMC data used in the scGEN tutorial^35^ to predict gene expression profiles of CD4+ T cells after interferon stimulation, given other cell types pre- and post-stimulation, and evaluated on the top 50 differentially expressed genes(Supplementary Methods).

We observed that which model performed best depended on which distance metric is used to evaluate the three models (Figure 3c). For example, R^2^, the metric used in the original scGen publication, reported that scGen with a shallow architecture vastly outperforms default scGen hyperparameters and also linear regression. However, evaluation with Pearson correlation provided exactly the opposite result. This makes it very difficult to decide which model better predicted gene expression during model development.

Based on its high CRP, robustness, and BioRep scores, we suggest using MSE and other type I distance metrics to evaluate gene expression predictions as a starting point to guide model development. In this case, MSE evaluation suggests that the shallow scGen model outperforms the default scGen and (marginally) the linear regression. We would also caution use of type III metrics in evaluating gene expression prediction, especially on high dimensional gene expression vectors. It is important to note that further downstream evaluation of a predictive model is still necessary to demonstrate a model’s goodness of fit. Nevertheless, we recommend that models which predict gene expression report MSE in their results.

### Use case 3: cell type response prioritization with distances

Another common task that relies on metrics is prioritization of cell types that respond most to perturbation. This task was formulated by Skinnider et al.^36^ who also introduced the Augur model to solve this task. We used the accompanying dataset from that study for this analysis. To demonstrate the utility of distances in data analysis, we used MSE to estimate cell type response magnitude as a proxy for response prioritization (Figure 3d) and compared the results to Augur. For a given dataset, Augur computes which cell type responds most to a given intervention and is methodologically similar to the classifier probability distance. Cell types that respond more are assigned a higher Augur score. We showed a positive correlation between Augur scores and MSE (Figure 3e) when MSE was used to measure the magnitude of cell type response, simply by taking the distance between the two conditions per cell type. We also showed that differential cell type prioritization using MSE recapitulates the differentially responding microglia and oligodendrocyte cell types exactly as in the example with Augur in Squair et al.^37^(Figure 3d). MSE provides a faster and more readily interpretable method of ranking cell types with comparable performance.

## Discussion

This work provides a quantitative comparison of distance metrics for single cell transcriptomics. Our observations and recommendations are broadly applicable whenever metric behavior is important, including for population comparison, functional analyses such as differential expression testing, and perturbation response prediction^19,36^. We find that conventional regression metrics such as MSE perform well on log-normalized scRNA-seq data, and, surprisingly, Spearman correlation does not, despite being a popular choice in the analysis of biological data^38^. The variety and number of datasets we tested allows us to summarize distance behavior into three types for end users. Finally, we show examples where the choice of distance metric can have a downstream impact. We conclude that MSE should be reported in model evaluation to ensure fair model comparability across studies and is a good starting point for investigating biological effect size and multi-condition analysis.

Despite consensus when it comes to fitting noise distributions, many non-NB and ZINB-parameterized scRNA-seq models, both those that predict gene expression and those that only model it, still perform well. General consensus suggests the Poisson, NB, and zero-inflated NB (ZINB) models as best capturing gene expression distributions^39–41^, with most scRNA simulators built on this insight^42^. While probabilistic models are popular^2,43–45^, optimal transport models are increasingly common^9,13,46^. Models such as scGen, which employs an MSE loss, perform competitively in batch correction and gene expression benchmarks^1,12,34^. The breadth of models and parameterizations in current literature suggests that, at present, there is no clear best architecture for gene expression prediction. Going forward, careful and robust model evaluation is key to advancement in the field^47^. In this work, we have built a stepping stone by recommending benchmarking and model evaluation measures, with modular distance calculations built on programmatically accessible datasets. We aim to move towards more standardized evaluation, which will only become increasingly necessary as better models develop and improvements become more incremental.

Our framework differs from prior work by evaluating distances between known intervention labels instead of cell type or batch labels^48,49^. While cell types are a common source of biological variation, they are typically manually annotated in single-cell data^20,50^. This manual annotation relies on dataset preprocessing and clustering algorithms which involve distance calculations^20^ and could bias results towards certain metrics. We initially chose intervention labels because it is commonly analyzed and a harder evaluation measure than cell type. The varying performance of distance metrics across datasets according to the BioRep metric (Figure 2b) suggests that the choice of metric might change depending on the biological covariate (e.g. cell type, timepoint, perturbation). This likely explains why our recommendation contradicts Huizing et al.^49^ recommendation of Wasserstein distance for batch and cell type distinguishability. Put another way, this challenges the assumption that all phenotype prediction can be assessed by the same measure, and by association, that all phenotypes can be modeled by the same distribution. This is supported by recent work demonstrating that the best parameters for negative binomial distributions are dataset-specific^48^, and that noise parameters change with time^51^.

While we aimed to provide a general distance metric recommendation, there are several limitations to our evaluation. There exist many other, less commonly used distances might have more optimal properties, including Mahalanobis distance, point cloud-based distances such as Chamfer and Hausdorff, and graph-based distances. Moreover, several metrics considered here benefit from being calculated in a different space than log-normalized data (Supplementary Figure 1d), and some distance measures are more commonly calculated on data that has been transformed into PCA space^19,33^. What effect additional processing^52^ has on observed differences, including the effect of gene selection methods, is a subject for future work.

Additionally, the availability of interventional datasets posed some limitations. The populations we considered are relatively homogeneous, with the exception of Santinha et al. where both control and perturbed existed in differentiated cell types in sufficiently high numbers. Other performance behavior might be uncovered with the additional analyses of datasets containing heterogeneous tissue models and perturbation.

In conclusion, we provide a general framework for assessing distance metrics in various datasets. Our reusable framework is publicly available at (https://github.com/theislab/perturbation-metrics) and users can efficiently call distance metric implementations from the pertpy package (https://github.com/theislab/pertpy) with an accompanying tutorial (https://pertpy.readthedocs.io/en/latest/tutorials/notebooks/distances.html). Any new metric can be easily assessed using this framework, facilitating the development of metrics for comparing transcriptomics distributions. Together with our metric recommendations, this framework adds clarity to robust analysis and model development for single-cell transcriptomics.

## Data and Code Availability

Distances were implemented in pertpy (https://pertpy.readthedocs.io). Reproducibility code and notebooks can be found at https://github.com/theislab/perturbation-metrics. All datasets used in this study are available from scPerturb^19^.

## Acknowledgements and Funding

We thank Niklas Schmacke, Linus Schumacher and Lukas Heumos for their valuable edits, comments, and recommendations on the manuscript. We thank Kemal Inecik for sharing his curation of Garcia 2022, and Lukas Heumos for helping integrate our software into the pertpy package. F.J.T. acknowledges support by the Helmholtz Association’s Initiative and Networking Fund through Helmholtz AI (ZT-I-PF-5-01) and by the European Union (ERC, DeepCell—101054957). L.Z. is funded by the Bavarian Ministry of Science and the Arts in the framework of the Bavarian Research Association ‘ForInter’ (Interaction of human brain cells). S.P. is supported by Open Targets (OTAR3083). C.S. acknowledges support by Wellcome-LEAP ΔTissue and the National Resource for Network Biology (NRNB) Research Resource (RR 031228-02). Views and opinions expressed are those of the author(s) only and do not necessarily reflect those of the European Union or the European Research Council. Neither the European Union nor the granting authority can be held responsible for them.

## Author contributions

Project was conceptualized by Y.J., T.D.G., S.P. and K.H. Formal analysis and investigation by Y.J. and M.L. Validation and visualization was performed by Y.J. Data was curated by T.D.G. and S.P. Methodology was developed by Y.J., T.D.G., and S.P. Extensive software contributions to the framework and package were made by Y.J., M.B., S.P., L.Z., and T.D.G. Funding acquisition, resources, and supervision were provided by C.S. and F.T. The original draft of this manuscript was written by Y.J., T.D.G., and S.P. All authors contributed to revisions.

