## Supplementary Figures and Methods for "Optimal distance metrics for single-cell RNA-seq populations"

### 1 Supplementary Figures and Tables

| First Author | Year | System (subset to) | Biological Covariate |
|---|---|---|---|
| Schiebinger | 2019 | 18-day MEF to iPSC reprogramming | post-reprogramming days |
| Garcia | 2022 | Ovarian interstitial cells | pcw |
| Norman | 2019 | leukemia cancer cell line (K562) | number of unique guides |
| Santinha | 2023 | AAV-Perturb-seq in mouse brain | N/A |
| McFarland | 2020 | esophageal carcinoma cell line (COLO680N) | time post drug treatment |
| Srivatsan | 2020 | breast cancer cell line (MCF7) | small molecule dose |
| Srivatsan | 2020 | leukemia cancer cell line (K562) | small molecule dose |
| Srivatsan | 2020 | lung cancer cell line (A549) | small molecule dose |

pcw: post-conception weeks

| First Author | Year | Description of Control | Number of Interventions* | Minimum Cells |
|---|---|---|---|---|
| Schiebinger | 2019 | day 1 of OSKM induction | 34 | 400 |
| Garcia | 2022 | 8.6 weeks post-conception | 13 | 300 |
| Norman | 2019 | non-targeting gRNA | 236 | 390 |
| Santinha | 2023 | safe-harbor gRNA | 29 | 480 |
| McFarland | 2020 | DMSO | 17 | 100 |
| Srivatsan | 2020 | DMSO | 752 | 270 |
| Srivatsan | 2020 | DMSO | 752 | 270 |
| Srivatsan | 2020 | DMSO | 752 | 270 |

*if applicable, number of conditions was subset to 100 for distance computations

Supplementary Table 1: Dataset information. Biological covariate was used for calculation of BioRep. Control condition was used for calculating CRP and robustness. Full citations are provided in the main text, and data is downloadable from scperturb.org.

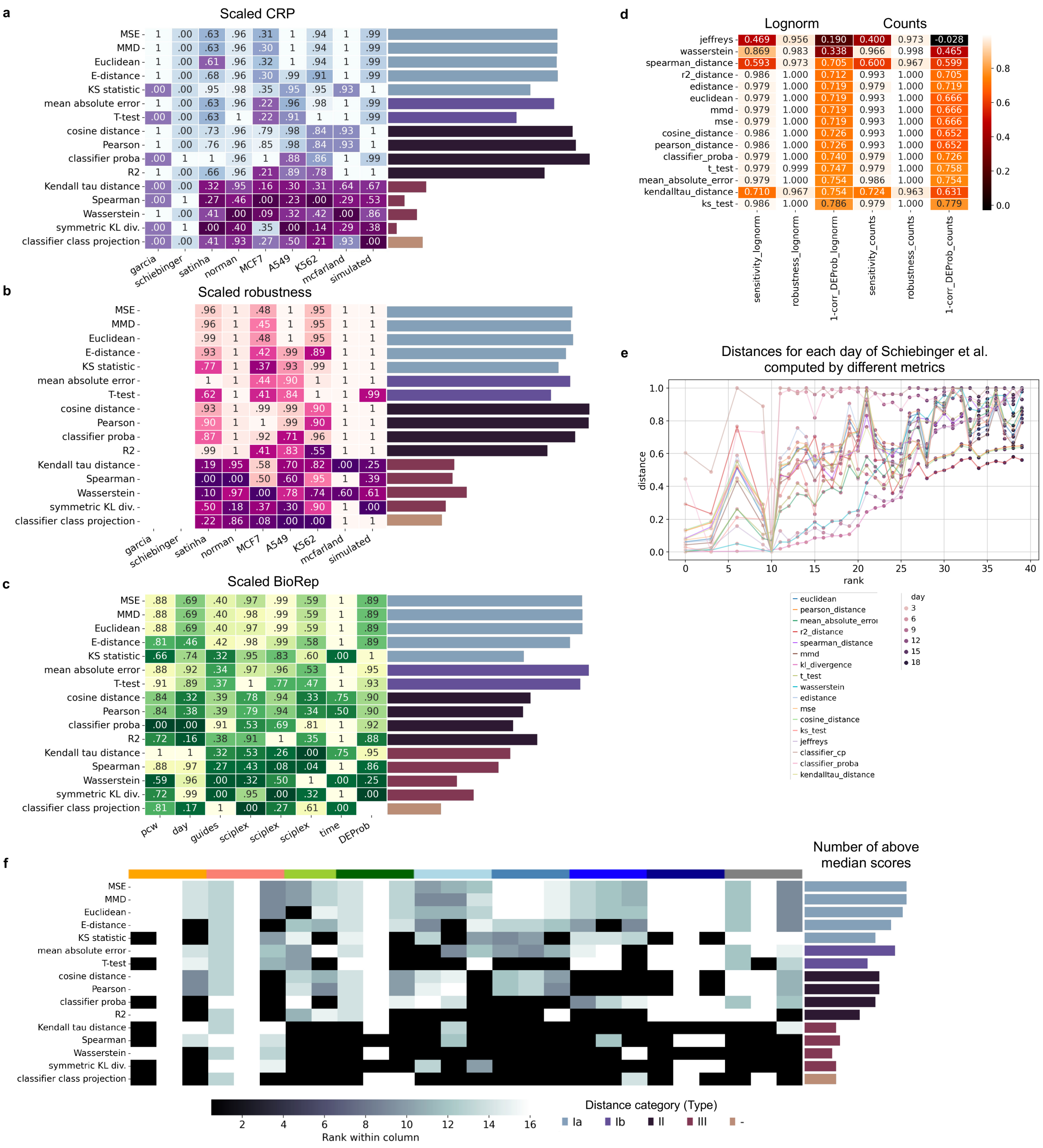

Supplementary Figure 1: In-depth look at evaluation results. Heatmaps of evaluation measure performance of (a) CRP, (b) robustness, and (c) BioRep. Each column is normalized between 0 and 1. The barplots on the right indicate the average score of each metric (row) for that task, and is colored by distance metric type from Table 1. Different types of metrics excel at different evaluation tasks. (d) Comparison of evaluating distances on log-normalized vs. raw count data using the simulated dataset. Relative values are extremely similar. (e) Distances from day 1 for each day of Schiebinger et al. Metrics with high BioRep score are those which increase monotonically with developmental time, most notably Wasserstein, which increases near linearly. (f) Heatmap of the results shown in Figure 2b. In (f), all scores worse than the median score of each column are colored black, showing that the top distance metrics performed consistently across all datasets. The number of non-black cells for each metric is shown visually in the bar plot on the right. The colored bar above the heatmap shows which columns came from different datasets, and correspond to the dataset colors and labels in Figure 2a.

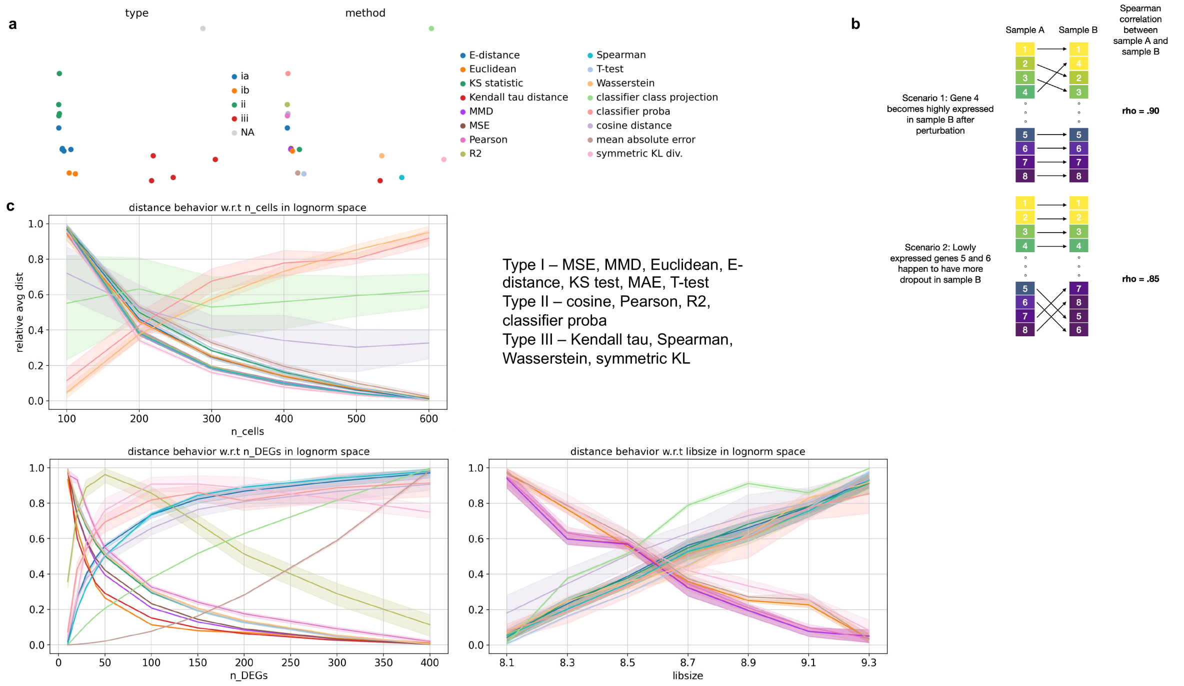

Supplementary Figure 2: Visual summaries of distances and their relationships. (a) PCA projection of distance scores across all tests, used to create the distance type categorizations. (b) Cartoon depiction of a situation in which Spearman and other type III metrics perform worse than other metric types. (c) Distance behaviors vary with respect to the number of cells, the number of differentially expressed genes (DEGs), and library size.

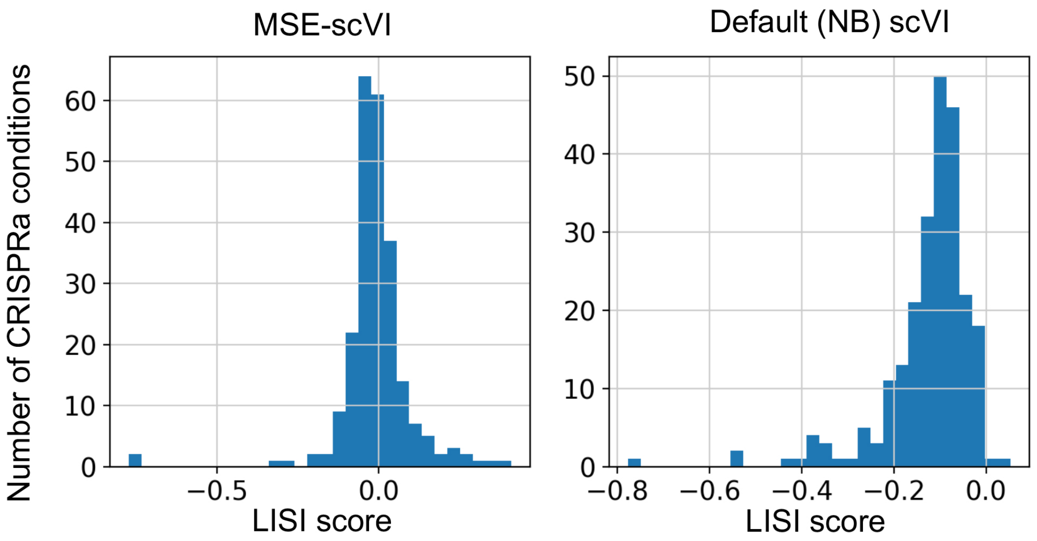

Supplementary Figure 3: Histograms of differential LISI scores of two different models. Left: MSE-scVI, which improves clustering of some CRISPRa perturbations but not others. Right: default scVI, which improves clustering of almost all CRISPRa perturbations.

### 2 Distance implementations

#### 2.1 Equations

For the purpose of the following definitions, we define $x^k$ as the gene expression vector of cell $k \in \{1, \ldots, N\}$. In addition, $x_i$ denotes the expression value of gene $i \in \{1, \ldots, n\}$, obtained by averaging expression over all cells in a sample. Put another way, equations which make use of $x_i$ or related terms use the centroids of each population. In cases in which the distance is not symmetric, $x$ denotes the perturbed conditions and $y$ denotes the control conditions. Unless noted otherwise, $\sum$ denotes $\sum_{i=0}^{n}$, the summation across all genes.

##### 2.1.1 Spearman's Rank distance

We rank genes by expression in each sample. Then,

$$\rho = \frac{6 \sum d_i^2}{n(n^2 - 1)} \tag{1}$$

Where $d_i$ represents the difference in rank of gene $i$ across both samples.

##### 2.1.2 Pearson's distance

$$r = 1 - \frac{\sum (x_i - \bar{x})(y_i - \bar{y})}{\sqrt{\sum (x_i - \bar{x})^2 \sum (y_i - \bar{y})^2}} \tag{2}$$

Where $\bar{x}$ and $\bar{y}$ are the means of $x$ and $y$ over all genes, respectively.

##### 2.1.3 R2 (Coefficient of Determination) distance

$$R^2 = \frac{\sum (x_i - y_i)^2}{\sum (x_i - \bar{x})^2} \tag{3}$$

Where $\bar{x}$ is the mean of the perturbed condition over all genes.

##### 2.1.4 Cosine distance

$$\text{Cosine distance} = 1 - \frac{\mathbf{x} \cdot \mathbf{y}}{|\mathbf{x}| \cdot |\mathbf{y}|} \tag{4}$$

Where $\mathbf{x}$ and $\mathbf{y}$ are gene expression vectors $[x_0, .., x_n]$ and $[y_0, .., y_n]$, and $\cdot$ represents the dot product.

##### 2.1.5 Wasserstein distance

$$W(p, q) = \inf_{\gamma \in \Gamma(p,q)} \int_{\mathcal{X} \times \mathcal{Y}} c(x, y)\, d\gamma(x, y) \tag{5}$$

Where $W(p, q)$ represents the first order Wasserstein distance between probability distributions $p$ and $q$, and $\Gamma(p, q)$ is the set of all joint distributions with marginals $p$ and $q$. The scalar-valued function $c(x, y)$ is the cost of transporting a unit of mass from $x$ to $y$, and $\mathcal{X}$ and $\mathcal{Y}$ are the support sets of $p$ and $q$, respectively. We used a Euclidean ground cost for $c$.

Empirical probability distributions are defined by sums of dirac delta functions representing the cells. In practice, we approximate the Wasserstein distance by the Sinkhorn distance, which solves a slightly modified, entropy-regularized version of the original optimization problem.

###### 2.1.6 Two-Sided T-statistic (Assuming Independent Samples)

$$t = \frac{1}{n} \sum \frac{x_i - y_i}{\sqrt{\frac{s_{x_i}^2}{n_x + \epsilon} + \frac{s_{y_i}^2}{n_y + \epsilon}}} \tag{6}$$

Where $s_{x_i}^2$ and $s_{y_i}^2$ are the variances of gene $i$ for perturbed and control, $n_x$ and $n_y$ are the sample sizes for perturbed and control, and $\epsilon$ is a small factor to avoid dividing by zero.

###### 2.1.7 Mean Absolute Error (MAE)

$$\text{MAE} = \frac{1}{n} \sum |x_i - y_i| \tag{7}$$

###### 2.1.8 Mean Squared Error (MSE)

$$\text{MSE} = \frac{1}{n} \sum (x_i - y_i)^2 \tag{8}$$

###### 2.1.9 Euclidean Distance

$$\text{Euclidean Distance} = \sqrt{\sum (x_i - y_i)^2} \tag{9}$$

###### 2.1.10 Kullback-Leibler (KL) Divergence

$$D_{KL}(P||Q) = \sum_{x \in \Omega} P(x) \log \left( \frac{P(x)}{Q(x)} \right) \tag{10}$$

Where $P, Q$ are two discrete probability distributions. The KL Divergence from $Q$ to $P$ is the expectation of the logarithmic difference between the probability distributions w.r.t. $P$.

In order to calculate KL divergence on non-discrete inputs, we use the formulation of KL divergence under the Gaussian assumption.

$$KL = \sum \ln \frac{s_{y_i}}{s_{x_i}} + \frac{s_{x_i}^2 + (x_i - y_i)^2}{2 * s_{y_i}^2} - \frac{1}{2} \tag{11}$$

Where $s$ is the standard deviation.

###### 2.1.11 Maximum Mean Discrepancy (MMD) with linear kernel

$$MMD^2 = \frac{1}{N(N-1)} \sum_{i=1}^{N} \sum_{j \neq i}^{N} k(x^i, x^j) - \frac{2}{NM} \sum_{i=1}^{N} \sum_{j=1}^{M} k(x^i, y^j) + \frac{1}{M(M-1)} \sum_{i=1}^{M} \sum_{j \neq i}^{M} k(y^i, y^j) \tag{12}$$

Where $n$ is the number of samples, $k(\cdot, \cdot)$ is the linear kernel function, and $\vec{x^i}$ and $\vec{y^i}$ are gene expression vectors for cells $i$ and $j$ respectively from two different samples.

###### 2.1.12 Kolmogorov–Smirnov test

For each gene $i$ we calculate the empirical distribution function $f_i(z) := |\{x_i^k : x_i^k \leq z, k = 1, \ldots, N\}|$ over all cells of the control condition and accordingly $\hat{f}_i$ for the perturbed condition. Per gene, the maximum distance between both distribution functions $\max_{z \geq 0} |f_i(z) - \hat{f}_i(z)|$ is recorded, and the results averaged over all genes to yield a single distance value.

###### 2.1.13 Kendall tau distance

Given two averaged gene expression vectors for two samples $x, y$, we count

- C: number of concordant pairs
- D: number of discordant pairs
- X: number of ties in x's ranking
- Y: number of ties in y's ranking

The Kendall tau distance is the defined as

$$\tau_{xy} = (C - D)/sqrt((C + D + X) * (C + D + Y)) \tag{13}$$

which we then normalized by

$$\tau'_{xy} = (1 - \tau_{xy}) \cdot \frac{n \cdot (n-1)}{4} \tag{14}$$

###### 2.1.14 Energy distance

We define

$$\delta_{XY} = \frac{1}{NM} \sum_{i=1}^{M} \sum_{j=1}^{N} ||x^i - y^j|| \tag{15}$$

$$\sigma_X = \frac{1}{N(1-N)} \sum_{i=1}^{N} \sum_{j=1}^{N} ||x^i - x^j|| \tag{16}$$

and $\sigma_Y$ accordingly. Then, the Energy distance is given by

$$E(X, Y) := 2\delta_{XY} - \sigma_X - \sigma_Y \tag{17}$$

###### 2.1.15 Classifier class projection

Given perturbations $P_i \in P_1, P_2...P_n$, control conditions $C_1, C_2...C_5$ and cells $x_i$ in $P_i$, we define the classifier class projection distance between $P_i$ and $C_i$ as follows. A linear regression classifier (*LogisticRegression* from the sklearn package[2]) was trained on all $x$ not in $P_i$ and all $C$. To compute the distance for the class $P_i$ and control subset $C_i$, we then retrieved the average post-softmax classification probabilities of all cells $\bar{x}_i$, and returned the probability of class $C_i$.

###### 2.1.16 Classifier control probability

Given perturbations $P$ and control condition $C$ we define the classifier class projection distance as follows. A linear regression classifier (*LogisticRegression* from the sklearn package[2]) was trained to distiguish $C$ and $P$ with 20 percent of $P$ held out. To compute the distance for the class $P_i$, we then retrieved the average post-softmax classification probabilities of all cells in $P_i$ and returned the probability of class $C$.

##### 2.2 Runtime

All metrics run within a few seconds for standard comparisons of one cell population against another. However, datasets with a large number of conditions or cells per condition, which are increasingly common, can take much longer. In particular, we note that Wasserstein and KS divergence become increasingly intractable on larger multi-condition datasets (Main Figure 3f). Generally, computation time for datasets increased linearly with respect to number of cells and conditions.

#### 3 Datasets

##### 3.1 Simulated data

Simulated data was generated using Splatter[4]. The goal of the simulation was to provide a dataset resembling small molecule perturbation in a cancer cell line, such as Srivatsan et al. or Norman et al. [3, 1]. As such, the control condition was modeled as a 20-step path with an endpoint that contained 0.05 DEProb in order to mimic natural variation in cell line data that might occur as a result of cell cycle or other effects.

32 perturbed conditions were generated using all pairwise combinations of eight values (DEProb, 0.00001, 0.00005, 0.0001, 0.0005, 0.001, 0.005, 0.01, 0.05) representing different probabilities that a gene is differentially expressed (DE) and four values (FacLoc,.2, .5, 1, 1.5) representing varying magnitudes of corresponding differential expression. 'FacScale', corresponding to variance of the distribution from which DE genes are drawn, was set equal to FacLoc. The probability of a gene increasing vs. decreasing after perturbation was equal (DownProb = 0.5). While this is not entirely realistic because perturbations generally have more upregulated genes than downregulated genes, we did not want to encounter artifacts of metric calculations due to this imbalance. Each perturbation was modeled as two steps. We simulated 40 thousand cells, with 40 percent of them as control.

We followed default parameters for the remaining simulation settings, with a few exceptions. Dropout was added with mid=0.3 in order to simulate additional noise. Library size was also decreased (lib.loc = 9) to simulate noisier conditions. Even still, and even with smaller DEProb, perturbations were generally easy to distinguish from control.

To assess the effect of library size on distance calculations using the simulation (Supplementary Figure 2c), we performed a subset of the above simulation five times but with different library sizes (8.1, 8.3, 8.5, 8.7, 8.9, 9.1, 9.3). Specifically, we only computed five perturbations instead of 32 for each of the library sizes to avoid long runtimes.

##### 3.2 Single-cell RNA-seq datasets

Before calculating distances, each dataset was filtered to contain one cell type (with the exception of Santinha et al.) and at most 100 conditions. The cell type was chosen in order to maximize the number of interventions with sufficient cell count. Interventions with fewer than the minimum number of cells listed in Supplementary Table 1 were removed. In general, we tried to use higher numbers of minimum cells were possible to give metrics a better chance at performing well. We also observed that metrics performed consistently past approximately 200 cells (Supplementary Figure 2). In order to avoid extreme runtimes, if more than 100 conditions remained, 100 conditions were randomly selected for distance computation. We then subset these conditions to the same number of cells (the number listed in Supplementary Table 1).

Then, each dataset was processed with the typical scanpy workflow (version 1.9.5). Cell-wise normalization to $10^6$ counts was applied, following which data was logarithmized. We computed 2000 highly variable genes using default parameters and computed all distances on this feature set.

To compute distances on raw count data, we followed the workflow above until the cell-wise normalization step. The same highly variable genes as calculated before were used to ensure any changes in distance calculations were due to differences between count data and log normalized data and not different features.

The following versions were used for relevant packages: anndata==0.8.0 umap==0.5.3 numpy==1.23.5 scipy==1.10.1 pandas==1.5.2 scikit-learn==1.2.2 statsmodels==0.14.0 igraph==0.10.8 pynndescent==0.5.8 pertpy==0.6.0

#### 4 Evaluation measures

For each distance metrics and dataset, we calculate the following three evaluation measures.

##### 4.1 Control rank percentile (CRP)

Intuitively, for each subsampled control population $c_j$, we compute the average distance $\bar{d}_{c_j}$ to the other control populations. We do the same for each perturbed population $p_i$ to get $\bar{d}_{p_i}$. This gives us a distance to control $d$ per $c$ and $p$, allowing us to compute the percentage of instances of $\bar{d}_p$ which is less than $\bar{d}_c$. One minus the average of this percentage across all $c_i$ is CRP. For the details of how the *distance* function in the algorithm above is computed, please see Supplementary section 2: Distance Implementations.

**Algorithm 1** Calculating control rank percentile for a single distance (CRP)

Let $C_1, \ldots, C_m$ be control subpopulations, $P_1, \ldots, P_n$ be different perturbations, $\bar{d_c} \leftarrow []$, and $\bar{d_p} \leftarrow []$.
**for** $i = 1 \rightarrow m$ **do**
  $d \leftarrow 0$
  **for** $j = 1 \rightarrow m$ **do**
    $d \leftarrow d + distance(C_i, C_j)$ if $C_i \neq C_j$
  **end for**
  $\bar{d}_{c_j} \leftarrow \frac{d}{m-1}$
**end for**
**for** $i = 1 \rightarrow n$ **do** /* Repeating for perturbed conditions. */
  $d \leftarrow 0$
  **for** $j = 1 \rightarrow m$ **do**
    $d \leftarrow d + distance(P_i, C_j)$
  **end for**
  $\bar{d}_{p_i} \leftarrow \frac{d}{n}$
**end for**
Let $r \leftarrow 0$ and $rank(x, y)$ be the rank of value $x$ in array $y$.
**for** $j = 1$ to $m$ **do**
  $r \leftarrow r + \frac{rank(\bar{d}_{c_j}, \bar{d}_p)}{n}$
**end for**
$CRP \leftarrow 1 - \frac{r}{m}$

##### 4.2 Robustness

Instead of averaging the above percentage, we take the inverse of the variance in this percentage as robustness.

##### 4.3 Calculation of differentially expressed genes

Differentially expressed genes was calculated between the control and stimulated condition in CD4+ T-cells only, using scanpy's $rank_genes_groups$ with default parameters.

#### 5 Methods related to distance metric use cases

##### 5.1 Modifying and training scVI with MSE loss

When scVI is set to model a negative binomial distribution, the posterior is parameterized by rate and scale parameters which are returned by two linear layers at the end of the decoder. In order to substitute in MSE loss to a fundamentally distribution-based model, we replaced the original reconstruction loss (the negative log likelihood of the negative binomial posterior w.r.t. $x$) with squared error loss of rate and $x$.

##### 5.2 Local inversion Simpson's Index (LISI)

LISI was computed using the 'lisi' function from the package scib-metrics. We computed LISI for every CRISPRa condition on the latent representation generated using scVI, MSE-scVI, or gene expression space (2000 highly variable genes). Supplementary Figure 3 was generated by subtracting, for each CRISPRa condition, the LISI score on gene expression space from the LISI score on latent representation space.

##### 5.3 Linear regression for comparison with scGen

For all models, we split the data into train and test where test contained CD4+ T-cells post stimulation. No stimulated CD4+ T-cells were seen in training.

Linear regression models are not made to predict completely unseen conditions as in scGen, but can do so nonetheless. We trained a *LinearRegression* from the sklearn package as follows: the control vs. stimulated label was encoded as a one-hot embedding and concatenated to the gene expression vector of the corresponding cell. At prediction time, the *LinearRegression* was provided with the gene expression vector of control cells, but the one-hot encoded vector representing stimulated. While this might seem dubious as the model never saw paired control and stimulated data, the performance was surprisingly and sufficiently good for our demonstration that evaluation metrics can differ in their reported outcomes.

All distance measures were calculated on the top 50 differentially expressed genes between stimulated CD4+ T-cells and unstimulated CD4+ T-cells.
